# Comparison of therapeutic effects between pulsed field ablation and cryoballoon ablation in the treatment of atrial fibrillation: a systematic review and meta-analysis

**DOI:** 10.1101/2024.04.24.591020

**Authors:** Yun Wan, Shuting Zeng, FuWei Liu, Xin Gao, Weidong Li, Kaifeng Liu, Jie He, Jianqing Ji, Jun Luo

## Abstract

**Background:** Pulmonary vein isolation (PVI) is the cornerstone of atrial fibrillation (AF) ablation surgery. Cryoballoon ablation (CBA), a conventional thermal ablation technique, enjoys widespread clinical application. In contrast, Pulsed field ablation (PFA) is a novel non thermal ablation technique for the treatment of atrial fibrillation (AF) patients, with safety comparable to traditional thermal ablation surgery. The present study aims to evaluate and compare the procedural efficiency and safety profiles of PFA and CBA in the management of AF.

**Methods:** We performed a systematic search across PubMed, the Cochrane Library, and Embase databases, encompassing the literature up to February 2024, to inform our systematic review and meta-analysis. When assessing outcome indicators, the risk ratio (RR) and its corresponding 95% confidence interval (CI) were calculated for dichotomous variables. For continuous variables, the mean difference (MD) and the associated 95% CI were determined. In this context, an RR less than 1 and an MD less than 0 were considered advantageous for the PFA group.

**Results:** In this analysis, nine observational studies encompassing 2,875 patients with AF were included. Among these, 38% (*n*=1105) were treated with PFA, while 62% (*n*=1,770) received CBA. The results indicated that PFA was associated with a significantly shorter surgical duration compared to CBA, with a mean difference (MD) of -10.49 minutes (95% CI [-15.50, -5.49]; *p*<0.0001). Additionally, the PFA group exhibited a reduced risk of perioperative complications relative to the CBA group, with a risk ratio (RR) of 0.52 (95% CI 0.30-0.89; *p*=0.02). Nevertheless, no statistically significant differences were observed when comparing the two treatment cohorts concerning fluorescence irradiation time (MD 0.71; 95% CI [-0.45, 1.86]; *p*=0.23) and the recurrence of atrial arrhythmias during follow-up (RR 0.95; 95% CI 0.78-1.14; *p*=0.57).

**Conclusions:** The outcomes of this investigation reveal that PFA holds a relative advantage over CBA in certain respects, notably by reducing both surgical duration and the incidence of perioperative complications. However, no significant distinction was identified between the two modalities concerning the duration of fluorescence irradiation or the rate of atrial arrhythmia recurrence. To enhance the robustness of these estimates, further research is needed, especially by incorporating additional randomized controlled trials.

## 1. Introduction

Atrial fibrillation (AF) is the most prevalent arrhythmia encountered in clinical practice, with its incidence and prevalence on a continual rise[1]. It can significantly diminish patients’ quality of life and elevate the risk of cardiovascular events[1, 2]. Pulmonary vein isolation (PVI) stands as the cornerstone procedure in all atrial fibrillation ablation surgeries and is strongly endorsed by current medical guidelines. The primary modalities employed, radiofrequency ablation (RFA) and low-temperature balloon cryoablation (CBA), have demonstrated both safety and efficacy[2-4]. However, as a conventional thermal ablation technique, CBA carries potential risks to adjacent structures of the pulmonary veins, including the phrenic nerve, esophagus, and pericardium[5-8].

Recently, pulsed field ablation (PFA) has emerged as a novel non-thermal ablation approach for achieving PVI in clinical settings[9]. PFA operates by applying a very brief high-voltage pulse field that generates an electric field, which disrupts the cell membrane by creating nanoscale pores, thereby inducing cell death[10-12]. A distinct advantage of PFA over traditional methods is its purported selectivity in inducing myocardial cell death, which theoretically minimizes damage to adjacent structures, including the esophagus and phrenic nerve, etc[9]. To date, several studies have attempted to compare the efficacy and safety of PFA with CBA in patients with atrial fibrillation, yet these studies are often constrained by limited sample sizes and inconsistent findings. To address this, we have conducted a systematic review and meta-analysis of pertinent literature, focusing on the comparative analysis of PFA and CBA with respect to surgical duration, fluoroscopy time, perioperative complications, and the recurrence of atrial arrhythmias.

## 2. Methods

This research protocol has been registered with the International Prospective Register of Systematic Reviews (PROSPERO ID: CRD42024513541). This study was designed in accordance with the PRISMA reporting guidelines for systematic evaluation and meta-analysis[13].

### 2.1 Search Strategy

As of February 2, 2024, two researchers, Y-Wan and ST Zeng, conducted independent searches for eligible studies using identical methodologies across the Cochrane Library, PubMed, and Embase databases. The search employed two sets of keywords, pertaining to “Pulsed field ablation” and “Cryoballoon ablation” which were combined using the logical operator “AND” to ensure a comprehensive search. The literature search was conducted without any language restrictions. Further details of the search strategy are outlined in Supplementary Table S2.

### 2.2 Eligibility criteria

Inclusion in this analysis requires that studies meet the following criteria: (1)Study Population: patients diagnosed with either paroxysmal or persistent atrial fibrillation.(2)Intervention and Comparison: Patients receiving PFA or CBA treatment.(3)Outcome Measures: The study should report on the efficacy of the surgical procedure, perioperative complications, or the recurrence of atrial arrhythmias.(4)Study Design: Eligible research designs include randomized and non-randomized controlled trials, case-control, cohort, and cross-sectional studies. Exclusion criteria are as follows: (1)Studies that are animal-based, reviews, case reports, letters, editorials, guidelines, systematic reviews, or conference abstracts are excluded.(2)Studies from which baseline data and the full text cannot be obtained are excluded.(3)In cases where a study has been reported multiple times, only the most recent results are considered.

### 2.3 Data extraction and quality assessment

Two researchers, Y-Wan and ST Zeng, independently conducted a comprehensive literature search, meticulously extracted data, and assessed the quality of each study. A third researcher, FW Liu, reviewed their work for accuracy and made the final determination on the inclusion of studies in the meta-analysis. The extraction process encompassed a range of details, including the first author’s name, year of publication, study design, geographical location of the study, characteristics of the study population, source of data, the manufacturer of the surgical instruments used, duration of follow-up, and both baseline and outcome data. To evaluate the quality of the observational studies included, we employed the improved Newcastle Ottawa Scale (NOS), a widely recognized tool that assigns a score ranging from 1 to 9 stars. This scale comprehensively assesses three critical aspects of study quality: selection of the study population, comparability between groups, and measurement of outcomes. A NOS score between 6 and 9 is indicative of high-quality research. A score within the range of 3 to 5 suggests research of satisfactory quality, while a score of 0 to 2 points to research of poor quality[14].

### 2.4 Statistical analysis

Summarize and analyze using Review Manager version 5.4(Cochrane Collaboration). For dichotomous outcomes, the relative risk (RR) was utilized as the effect measure, while the mean difference (MD), derived from the inverse variance, was applied for continuous outcomes. In the context of this analysis, an RR value less than 1 and an MD value less than 0 were considered advantageous for the PFA group. To assess the variability among studies, the I^2^ statistic and the Cochran Q-test were employed. Heterogeneity was deemed significant when the I^2^ statistic exceeded 50% and the p-value was less than 0.1. Under such circumstances, a random effects model was engaged for the analysis. In cases where heterogeneity was not significant, a fixed effects model was utilized. To mitigate the influence of inappropriate interference, sensitivity analyses were conducted. Each study was sequentially omitted, prompting a recalculation of the pooled effects and heterogeneity. A study was deemed influential if its removal accounted for the heterogeneity or altered the significance of the effect[15]. Given the limited number of studies included, falling below the threshold of ten, a publication bias analysis was not performed[16]. A p-value less than 0.05 was established as the threshold for statistical significance.

## 3. Results

### 3.1 Study selection

A comprehensive search of PubMed, Embase, and Cochrane electronic databases yielded a total of 118 studies, following a pre-planned retrieval strategy. After the removal of 27 duplicates, 91 studies remained. Upon reviewing the titles and abstracts, 75 studies were excluded. The full texts of the remaining 16 articles were then assessed, resulting in the exclusion of 7 articles for the following reasons: (1) lack of relevant data (*n*=4); (2) Lack of control group (*n*=1); (3) Unable to obtain full text (*n*=1); (4) The target audience does not match (*n*=1). The specific reasons for excluding each study are detailed in Supplementary Table S3. Ultimately, we included a total of 9 studies (Figure 1), comprising 6 cohort studies[17-22] and 3 case-control studies[23-25] .

**Figure1.**
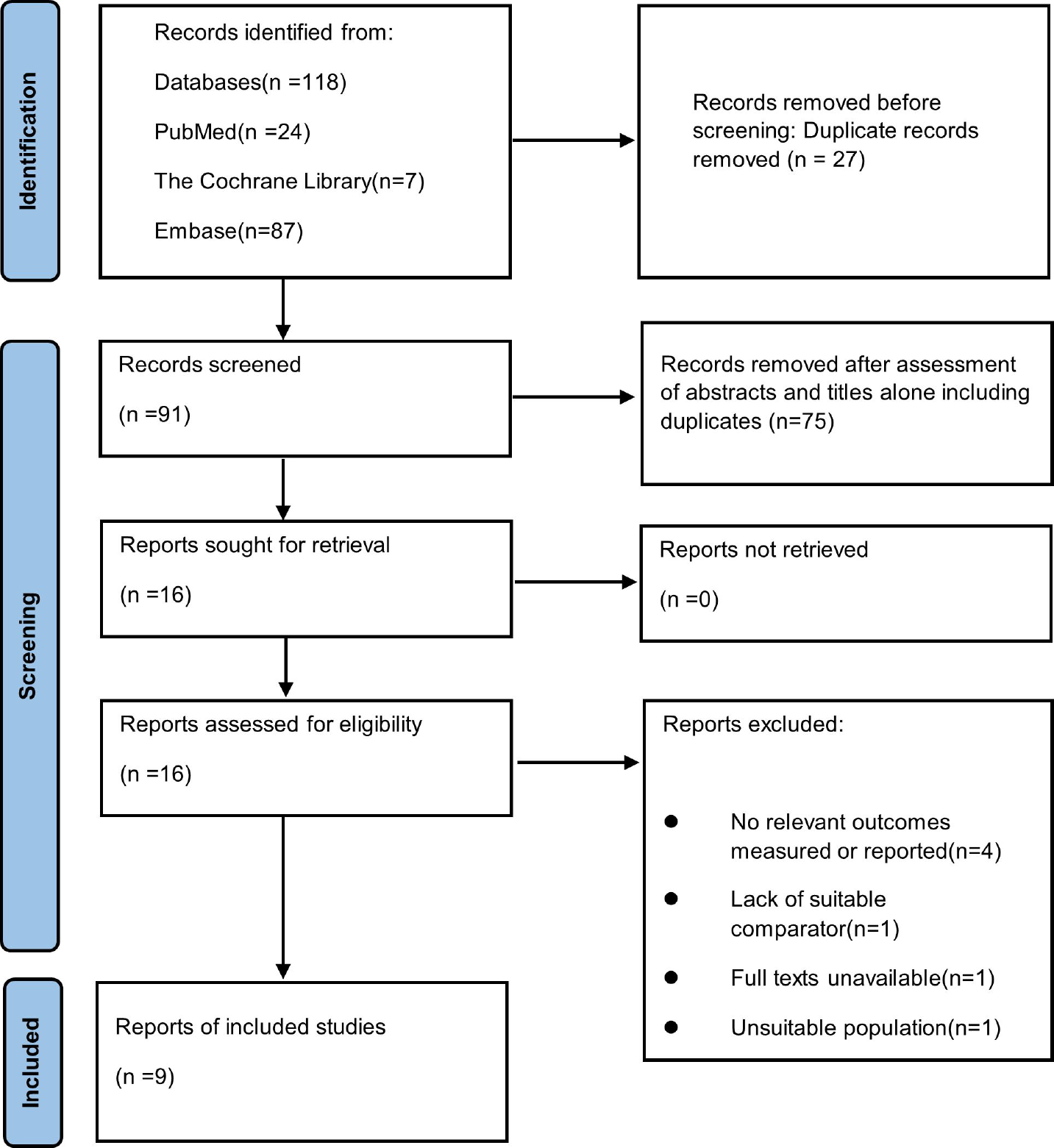
PRISMA Flow diagram of included studies.

### 3.2 Study characteristics and quality assessment

Table 1 encapsulates the baseline characteristics of the nine included studies, encompassing a total of 2,875 patients, with 65% being male. Specifically, 1105 cases (38%) were treated using PFA, while 1770 cases (62%) received CBA. The sample size across the studies varied from 43 to 1714 patients, with an average age ranging from 39.9 to 68.9 years. The research features are shown in Table 2. One of them is from Switzerland[18], one is from Netherlands[21], and the others are all from Germany. Five studies included patients with atrial fibrillation who underwent ablation surgery for the first time. To rigorously assess the quality of the included observational studies, the Newcastle-Ottawa Scale (NOS) was employed. This scale was used to evaluate the quality of the six cohort studies and three case-control studies, respectively. Gratifyingly, all studies were deemed high-quality, with NOS scores exceeding six stars. For a detailed breakdown of the quality assessment, refer to Supplementary Table S4-5.

**Table 1.** Baseline characteristics of included studies.

| study ID | group | Patients<br>(*n*) | Age (years) | Male,<br>*n*(%) | BMI<br>(kg/m<sup>2</sup>) | Persistent<br>AF, *n* (%) | Congestive<br>heart<br>failure, *n*<br>(%) | Hypertension,<br>*n* (%) | Diabetes mellitus,<br>*n* (%) | Stroke/TIA,<br>*n* (%) | CAD/vascular<br>disease, *n* (%) | Dyslipidemia,<br>*n* (%) | CHA<sub>2</sub>DS<sub>2</sub>-VASc<br>score | Ejection<br>fraction<br>(%) | Left atrial<br>diameter<br>(mm) | Class<br>I/III<br>AADs,<br>*n*(%) |
| --- | --- | --- | --- | --- | --- | --- | --- | --- | --- | --- | --- | --- | --- | --- | --- | --- |
| Schipper, J. | PFA | 54 | 69±11 | 37(68.52) | 27.8±5 | 38(70.37) | NR | 39(72.22) | 9(16.67) | NR | 17(31.48) | 19(35.19) | 3.0±1.8 | 53.3±10.9 | 38.8±5.8 | 18(33.33) |
| H-2023 | CB | 54 | 67±13 | 37(68.52) | 28.1±4.5 | 37(68.52) | NR | 37(68.52) | 9(16.67) | NR | 14(25.93) | 18(33.33) | 2.7±1.7 | 54.9±10.6 | 39.6±6.1 | 15(27.78) |
| Badertscher, | PFA | 106 | 64.9±9.42 | 67(63.21) | 27.35±3.76 | 41(38.68) | NR | 64(60.38) | 12(11.32) | NR | 9(8.49) | 45(42.45) | NR | 57±8.6 | 41±6.4 | NR |
| P-2023 | CB | 75 | 63.5±9.97 | 48(64) | 26.65±3.78 | 24(32) | NR | 35(46.67) | 6(8) | NR | 6(8) | 20(26.67) | NR | 58±9.1 | 40±6.3 | NR |
| Tohoku, | PFA | 54 | 69±9 | 33(61.11) | 28±5 | 22(40.74) | 5(9.26) | 37(68.52) | 7(12.96) | 2(3.70) | 3(5.56) | NR | NR | 60.8±7.2 | 40.5±5.9 | 9(16.67) |
| S-2023 | CB | 43 | 69±9 | 19(44.19) | 28±6 | 18(41.86) | 4(9.30) | 30(69.77) | 2(4.65) | 4(9.30) | 8(18.60) | NR | NR | 59.1±13.4 | 39.5±7 | 9(20.93) |
| Lemoine, | PFA | 51 | 68±12 | 36(70.59) | 27±5 | 30(58.82) | 14(27.45) | 34(66.67) | 8(15.69) | 1(1.96) | 16(31.37) | NR | 2.7±1.7 | 52±12 | 37±14<sup>a</sup> | 13(25.49) |
| M. D-2023 | CB | 40 | 63±13 | 26(65) | 28±5 | 21(52.5) | 7(17.5) | 25(62.5) | 1(2.5) | 3(7.5) | 7(17.5) | NR | 2.3±1.6 | 53±9 | 36±13<sup>a</sup> | 5(12.5) |
| Rattka, | PFA | 94 | 63±12 | 58(61.72) | NR | 28(29.79) | NR | 73(77.66) | 16(17.02) | 7(7.45) | 33(35.11) | 59(62.77) | 2.65±2.26 | 54.30±9.03 | NR | 28(29.79) |
| M-2024 | CB | 47 | 64±12 | 35(74.47) | NR | 18(38.30) | NR | 26(55.32) | 15(31.91) | 2(4.26) | 21(44.68) | 32(68.09) | 2.65±2.29 | 56.06±3.82 | NR | 16(34.04) |
| Blockhaus, | PFA | 23 | 57.13±10.32 | 15(65.22) | 28.13±3.6 | 11(47.82) | 3(13.04) | 15(65.22) | NR | 0 | 2(8.70) | NR | 1.52±1.12 | 55.65±7.6 | 41.22±3.14 | 3(13.04) |
| C-2023 | CB | 20 | 59.1±8.96 | 16(80) | 26.27±3.78 | 10(50) | 5(25) | 8(40) | NR | 2(10) | 5(25) | NR | 1.65±1.35 | 55±8.11 | 41±2.77 | 4(20) |
| van de Kar, | PFA | 473 | 64.65±9.67 | 301(63.64) | 26.91±3.79 | 182(38.48) | NR | NR | 44(9.3) | NR | NR | NR | NR | 55 | 35±10.42<sup>a</sup> | NR |
| M. R.<br>D-2024 | CB | 1241 | 63.65±9.65 | 859(69.22) | 27.05±3.79 | 422(34.00) | NR | NR | 90(7.25) | NR | NR | NR | NR | 55 | 36.35±11.14<sup>a</sup> | NR |
| Urbanek, | PFA | 200 | 69.95±11.20 | 118(59) | 27.35±5.23 | 84(42) | 27(13.5) | 132(66) | 28(14) | 10(5) | 28(14) | NR | 2.70±1.49 | NR | 41.35±6.72 | NR |
| L-2023 | CB | 200 | 67.65±14.19 | 108(54) | 27.70±4.48 | 73(36.5) | 23(11.5) | 141(70.5) | 33(16.5) | 15(7.5) | 27(13.5) | NR | 2.65±2.24 | NR | 40±5.97 | NR |
| Wahedi, | PFA | 50 | 67±10.6 | 32(64) | 26.2±3.5 | 14(28) | 5(10) | 30(60) | 3(6) | 2(4) | 9(18) | NR | 2.2±1.5 | NR | 41.5±6.3 | 15(30%)<sup>b</sup> |
| R-2024 | CB | 50 | 65±10.6 | 29(58) | 29.4±8.6 | 29(58) | 5(10) | 35(70) | 8(16) | 3(6) | 11(22) | NR | 2.4±1.3 | NR | 42.5±7.6 | 12(24)<sup>b</sup> |

Abbreviations: BMI, body mass index; TIA, transient ischemic attack; CAD, coronary artery disease; AAD, anti-arrhythmic drug; PFA, Pulsed field ablation; CB, Cryoballoon ablation
<sup>a</sup> Left Atrial Volume index; <sup>b</sup> Only taking AADs

**Table 2.** Study characteristics of included studies.

| study | Study design | Country | Study population | Pulsed field ablation catheter | Cryoballoon Ablation catheter | Follow-up | Monitoring |
| --- | --- | --- | --- | --- | --- | --- | --- |
| Schipper, J. H-2023 | retrospective,single-center,non-randomized,controlled trial | Germany | Patients with paroxysmal or persistent atrial fibrillation undergoing PVI for the first time | FARAPULSE™ system (Boston Scientific) | The 31mm POLARx™ cryoablation system (Boston Scientific) | 273±129 days | 12-lead ECG and 24-h Holter at 3, 12 months |
| Badertscher, P-2023 | Prospective,single-center,non-randomized,controlled trial | Switzerland | Paroxysmal or persistent atrial fibrillation patients aged 18 and above who receive PVI for the first time | FARAPULSE™ system (Boston Scientific) | The POLARx™ Cryoablation System (Boston Scientific) | 390±265 days | 12 - lead ECG, and 7-d Holter at 3, 6, and 12 months |
| Tohoku, S-2023 | Prospective,single-center,non-randomized,controlled trial | Germany | Symptomatic drug-resistant atrial fibrillation patients aged 18-85 | FARAPULSE™ system (Boston Scientific) | Flexcath, Medtronic | PFA group followed up for 328±114 days | 24-h Holter at 3, 6 months, if symptomatic |
| Lemoine, M. D-2023 | case-control study | Germany | Cardiac arrhythmia patients over 18 years old | FARAPULSE™ system (Boston Scientific) | Arctic Front Advance, Medtronic | NR | NR |
| Rattka, M-2024 | retrospective,single-center,non-randomized,controlled trial | Germany | Patients with paroxysmal or persistent atrial fibrillation undergoing PVI for the first time | FARAPULSE™ system (Boston Scientific) | The POLARx™ Cryoablation System (Boston Scientific) | 365 days | Tlinical assessment, TTE and 7-d Holter at 3, 6, and 12 months |
| Blockhaus, C-2023 | case-control study | Germany | Patients with paroxysmal or persistent atrial fibrillation | Lasso, Biosense Webster | 2nd generation, Arctic Front Advance, Medtronic | NR | NR |
| van de Kar, M. R. D-2024 | retrospective,single-center,non-randomized,controlled trial | Netherlands | Paroxysmal or persistent atrial fibrillation patients aged 18 and above who receive PVI for the first time | FARAPULSE™ system (Boston Scientific) | The 27-mm Arctic Front™ catheter, Medtronic | 180 days | Assess symptoms at 6 months |
| Urbanek, L-2023 | retrospective,single-center,non-randomized,controlled trial | Germany | Symptomatic atrial fibrillation (paroxysmal and persistent atrial fibrillation) | FARAPULSE™ system (Boston Scientific) | CB2, 28-mm Arctic Front Advance, Medtronic | 383±28 days( PFA group);<br>364±241 days( CBA group) | 72-h Holter at 6, 12 months |
| Wahedi, R-2024 | case-control study | Germany | Paroxysmal or persistent atrial fibrillation patients undergoing PVI for the first-time using PFA or CB | FARAPULSE™ system (Boston Scientific) | fourth-generation Arctic Front Advance, Medtronic | NR | NR |

### 3.3 Outcomes of the included studies

#### 3.3.1 Procedure duration

In different studies, there is a significant difference in surgical time. For PFA, the average surgical time ranges from 34.5-162 minutes. Estimate the summary effects through a random effects model. The results showed that PFA had a shorter surgical time compared to CBA (MD -10.49; 95% CI [-15.50, -5.49]; *p*<0.0001). However, the heterogeneity between studies was significant (*p*<0.00001, I^2^= 90%) (Figure 2-A).

**Figure2.**
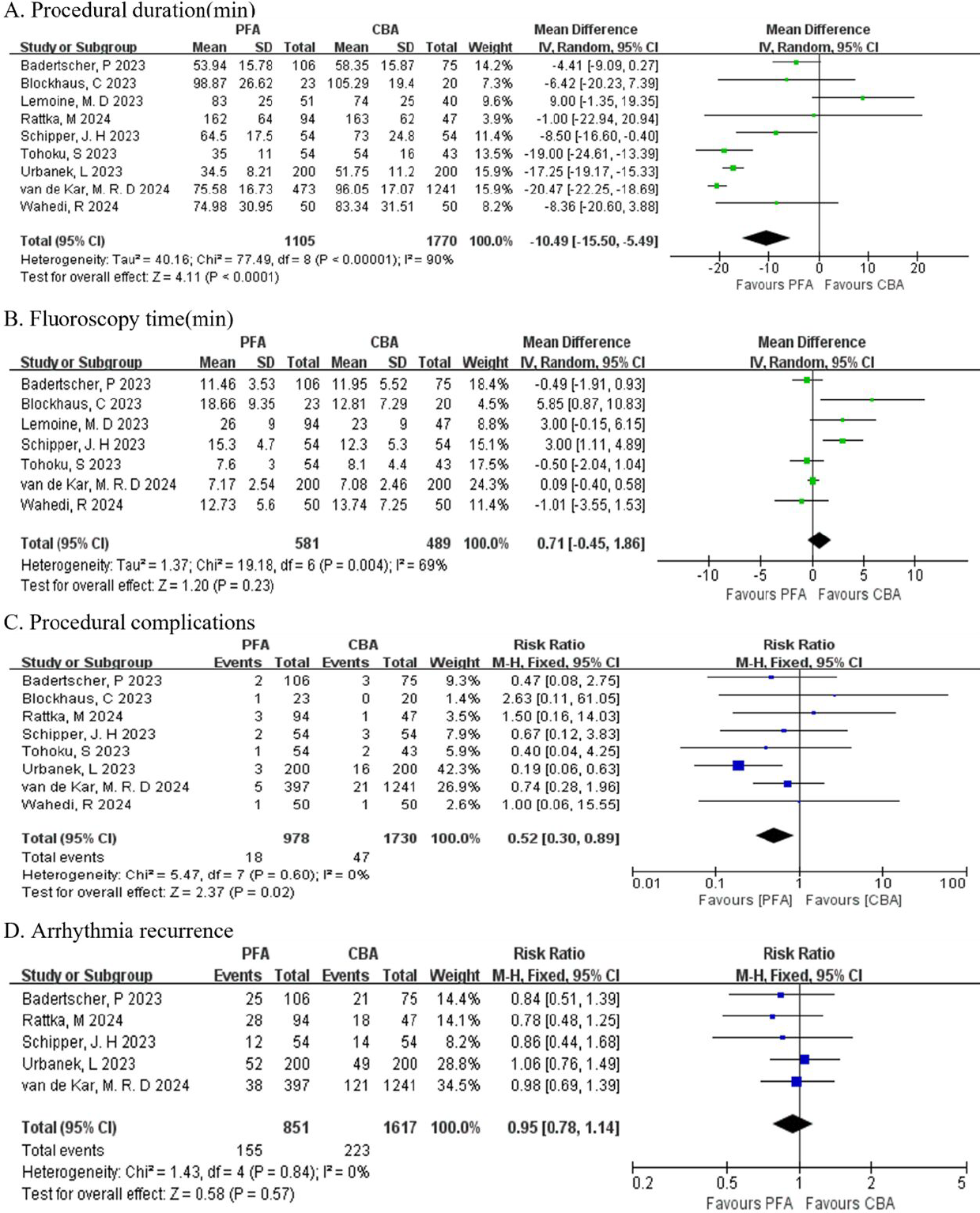
Forest plot comparison of pulse field ablation and cryoballoon ablation in perioperative and postoperative follow-up of patients with atrial fibrillation. A Comparison of procedure duration. B Comparison of fluoroscopy time. C Comparison of procedural complications. D Comparison of arrhythmia recurrence during follow-up period.

#### 3.3.2 Fluoroscopy time

In the seven studies analyzed, the duration of fluorescence irradiation for the PFA group varied, with a range from 7.17 to 26 minutes. For the CBA group, the corresponding range was 7.08 to 23 minutes. A comparison of the two groups revealed no statistically significant difference in the fluorescence irradiation time between PFA and CBA (MD 0.71; 95% CI, [-0.45, 1.86]; *p*=0.23). However, the heterogeneity between studies is significant (*p*=0.004, I ^2^= 69%) (Figure 2-B).

#### 3.3.3 Complications

The perioperative complications observed in different studies are not consistent. Notably, only the study by Schipper, J. H. reported on the occurrence of perioperative pulmonary vein stenosis, with no cases observed in either the PFA or CBA groups. In the studies conducted by Schipper, J. H., Wahedi, R., and Urbanek, L., the incidence of atrial esophageal fistula was exceptionally rare, with a single case noted during the follow-up in Urbanek, L.’s research. The patient in question experienced spontaneous improvement without the need for further intervention. Lemoine, M. D.’s study did not report any perioperative complications. The remaining eight studies provided a comprehensive comparison of perioperative complications between the two ablation groups, encompassing diaphragmatic nerve paralysis, pericardial tamponade, and stroke/transient ischemic attack(TIA).The risk of perioperative complications for the PFA group was found to be lower than that for the CBA group, with a relative risk (RR) of 0.52 (95% CI 0.30-0.89; *p*=0.02). The heterogeneity between these studies was minimal (*p*=0.60, I^2^=0%)(Figure 2-C).

#### 3.3.4 Recurrence of AF

Among the six studies that provided follow-up information, one reported a follow-up duration of six months, whereas the other five studies had an average follow-up period of twelve months. Analysis of postoperative follow-up data revealed no statistically significant difference in the recurrence of atrial arrhythmias between the PFA group and the CBA group (RR 0.95; 95% CI 0.78-1.14; *p*=0.57). Heterogeneity among these studies was extremely low (*p*=0.84, I^2^=0%)(Figure 2-D).

### 3.4 Sensitivity analysis

We performed a sensitivity analysis to address the significant heterogeneity observed in surgical time and fluorescence irradiation time across various studies. The stability of the pooled results was maintained when each study was sequentially omitted. For surgical time, the mean difference (MD) ranged from -12.95 to -8.18, with an I^2^ statistic between 85% and 91%. In the case of fluorescence irradiation time, the MD ranged from 0.16 to 1.09, and the I ^2^ statistic varied from 52% to 73%. The heterogeneity findings were consistent with the overall combined analysis, confirming the stability of our conclusions (see Supplementary Tables S6-S7 for details).

## 4. Discussion

In recent years, the use of PFA in patients with AF has been on the rise. A series of single-arm clinical trials have affirmed the clinical efficacy and safety of PFA in the treatment of AF patients[9, 26-28]. The effectiveness of ablation surgery is gauged by parameters such as the success rate of Pulmonary Vein Isolation (PVI), the duration of the surgical procedure, the time spent on fluorescence irradiation, and the rate of atrial arrhythmia recurrence. The safety profile is assessed based on the incidence of peri-operative complications and adverse events. To compare the efficacy and safety of PFA compared to CBA in patients with AF, we conducted this systematic review and meta-analysis. Our study encompassed nine clinical studies, comprising six single-center cohort studies and three case-control studies. Notably, the study led by van de Kar, M.R. et al., a retrospective analysis from a high-volume center in the Netherlands, stands out for its large patient cohort, involving 1714 participants. We will delve into the following outcome measures for our analysis.

### 4.1 Successful Pulmonary Vein Isolation (PVI)

PVI is one of the main determining factors influencing the effectiveness of PFA[9]. Among the nine studies included in our analysis, seven provided data on the success rate of PVI. In the study by Tohoku et al., the success rate was notably 88.69% for the CBA group and a perfect 100% for the PFA group. This discrepancy is not explained in this paper, and we believe that it may be caused by insufficient sample size or improper operation of surgical personnel. Success rates in other studies ranged from 99% to 100%. Given the minor differences observed, we did not conduct further detailed summary or analysis of these data. Collectively, these findings suggest that both PFA and CBA are effective in achieving successful ablation of lesions in patients with Atrial Fibrillation (AF).

### 4.2 Procedural Characteristics

The findings from this meta-analysis indicate that PFA reduces the duration of surgical procedures when compared to CBA. No significant disparity was observed in the duration of fluorescence fluoroscopy between the two methodologies. This underscores the efficacy of PFA in the treatment of patients with Atrial Fibrillation (AF). However, considerable variability and high heterogeneity were noted in the surgical times reported across different studies, a discrepancy that could be attributed to the lack of a uniform definition of surgical time. For instance, some studies have adopted the “skin to skin” approach, defining surgical time as the interval commencing from inguinal puncture and concluding with the sheath removal[17, 18, 24] . Conversely, other studies have measured surgical time from the moment of entry into the operating room to the time of departure[21]. A subset of studies did not provide a detailed explanation of their definition of surgical time. Among the studies included in this analysis, Badertscher, P et al. documented the left atrial residence time, which was 38 (30-49) minutes for PFA and 37 (31-48) minutes for CBA. This parameter of left atrial residence time is posited as a more accurate reflection of the surgical time. The reason for this is that additional procedures, such as the insertion and removal of sheaths, which are not accounted for in the “skin-to-skin time” measurement, may introduce a more pronounced variation in the reported surgical times[29]. Furthermore, other factors contribute to the observed heterogeneity among studies. These factors include differences in the technical levels of the surgical teams performing the procedures, variations in the instruments utilized, and the distinct anatomical positioning of the pulmonary veins. The variability in these aspects can introduce significant differences in the surgical outcomes and times reported in the literature.

### 4.3 Adverse Events/Outcomes

At present, some studies have reported common adverse events of PFA, including diaphragmatic nerve paralysis, surgical site bleeding, pericardial tamponade, cerebral embolism, cardiac conduction block, pericarditis, death, atrial esophageal fistula, and pulmonary vein stenosis[30-34]. It is important to note that the perioperative complications noted by various research institutes are not uniform, leading to a decision not to provide a generalized summary of complication rates across studies. One of the studies only observed one complication of Stroke/TIA[23], and there was no occurrence in the experimental and control groups. Some studies have observed esophageal injury, with only one case observed in the CBA group[22], and the lesion naturally healed without additional treatment. One study observed pulmonary vein stenosis^[17]^, and no occurrence was observed in both groups. Although surgical induced pulmonary vein stenosis was not reported in the included studies, the sample size was small and insufficient to detect significant differences. According to available data, the risk of severe pulmonary vein stenosis as a result of atrial fibrillation ablation procedures ranges from 0.32% to 3.4%[35]. To ascertain the true incidence of this complication, there is a clear need for larger-scale studies. Regarding the insertion site for the sheath, the inguinal area is commonly selected for puncture and sheath insertion, followed by entry into the left atrium via atrial septal puncture. During this phase, the majority of observed complications were related to vascular issues such as pseudoaneurysms, arteriovenous fistulas, and atrioventricular fistulas. Statistical analysis revealed no significant difference in the surgical incidence between the two groups.

In comparison to other complications, diaphragmatic nerve paralysis, pericardial tamponade, and Stroke/TIA have each been documented in eight separate studies. Specifically, for diaphragmatic nerve paralysis, van de Kar, M. R. D et al. reported that 15 individuals (1.2%) in the CBA group experienced this condition, with 9 of these cases being transient. In the study conducted by Urbanek, L et al., the prevalence of diaphragmatic nerve paralysis was notably higher in the CBA group, reaching 7.5%, predominantly transient in nature. After comparing various studies, it was found that PFA is safer than CBA in terms of the incidence of perioperative phrenic nerve injury (RR 0.15; 95% CI 0.06-0.39; *p*<0.0001). When examining pericardial tamponade, a common consequence of cardiac perforation observed in some studies, the incidents can often be attributed to unintended muscle or diaphragmatic convulsions[36, 37] . A comparative analysis of the studies revealed that the PFA group faced a higher risk of pericardial tamponade relative to the CBA group (RR 3.48; 95% CI 1.26-9.66; *p*=0.02). Most instances of pericardial tamponade can be resolved with pericardial puncture drainage within 24 hours, and there was one case that improved following surgical intervention[17]. In the context of Stroke/TIA, no significant difference was observed between the PFA and CBA groups (RR 0.99; 95% CI 0.30-3.22; *p*=0.99). After evaluating the collective risks associated with diaphragmatic nerve paralysis, pericardial tamponade, and Stroke/TIA, it was concluded that the PFA group exhibited a lower risk of perioperative complications compared to the CBA group (RR 0.52; 95% CI 0.30-0.89; *p*=0.02). Combining PFA has theoretical safety advantages, and this study further confirms its safety in clinical applications.

### 4.4 Recurrence of AF or Other Atrial Arrhythmias

Among the six cohort studies included, one study only followed up on the PFA group[19]. The remaining five studies provided follow-up data for both the PFA and CBA groups, predominantly with a follow-up duration of 12 months. The primary endpoint of follow-up in these studies is the recurrence of atrial fibrillation or other atrial arrhythmias, with a defined blank period of 3 months post-surgery completion. Upon examining the recurrence rates reported across these studies, no significant disparity was identified between the PFA and CBA groups (RR 0.95; 95% CI 0.78-1.14; *p*=0.57). It is worth highlighting a particular study by van de Kar, M. R. D, which had a follow-up period of 6 months. In this study, the rates of re-ablation were reported, with 9.57% for the PFA group and 9.75% for the CBA group.

## 5. Limitations

This study is subject to several limitations that warrant consideration. Firstly, the pool of studies included in this analysis is relatively limited, and all are single-center, non-randomized, and observational in nature. Given the inherent constraints of observational research, the quality of evidence presented is not optimal. Some of the included studies have a limited scope in observing perioperative complications, which may impact the comparative safety assessment between PFA and CBA. Furthermore, most studies have a limited number of participants, and the cohorts in each study include a mix of patients with paroxysmal atrial fibrillation and persistent atrial fibrillation. This diversity within the study populations prevents us from conducting sub-group analyses based on the type of atrial fibrillation. Lastly, variations in the expertise and experience of the surgical teams across the different studies may also introduce bias and impact the study outcomes. In light of these limitations, it is imperative that further research be conducted to validate and reinforce the findings of this analysis. The pursuit of more extensive, multicenter, randomized, and controlled studies would be beneficial in providing a more robust and reliable evidence base for the comparison of PFA and CBA in the treatment of atrial fibrillation.

## 6. Conclusions

Drawing from the findings of this meta-analysis, it appears that PFA for the treatment of atrial fibrillation can reduce surgical time and is associated with a lower risk of surgical complications when compared to CBA. Additionally, no significant differences were observed between the two methods in terms of fluorescence irradiation time and the recurrence rate of atrial arrhythmias. However, to more definitively establish the comparative effectiveness and safety profiles of PFA and CBA, more high-quality research is needed to provide evidence.

## Data availability

The data of this study are detailed within the article itself and are also available in the Supplementary Material. For any additional inquiries or questions, please reach out to the corresponding authors.

## Additional file

**Table S1**. PRISMA Checklist. **Table S2**. The search strategies of this meta-analysis. **Table S3**. Studies excluded (n=7) with reasons. **Table S4**. Quality assessment of the included Cohort studies by Newcastle–Ottawa scale. **Table S5**. Quality assessment of the included Case-control studies by Newcastle–Ottawa scale. **Table S6**. The heterogeneity of the included studies through sensitivity analysis - Procedure duration. **Table S7**. The heterogeneity of the included studies through sensitivity analysis - Fluoroscopy time.

